# Computational characteristics of interictal EEG as objective markers of epileptic spasms

**DOI:** 10.1101/2020.11.13.380691

**Authors:** Rachel J. Smith, Derek K. Hu, Daniel W. Shrey, Rajsekar Rajaraman, Shaun A. Hussain, Beth A. Lopour

## Abstract

**Objective:** Favorable neurodevelopmental outcomes in epileptic spasms (ES) are tied to early diagnosis and prompt treatment, but uncertainty in the identification of the disease can delay this process. Therefore, we investigated five computational electroencephalographic (EEG) measures as markers of ES.

**Methods:** We measured 1) amplitude, 2) power spectra, 3) entropy, 4) long-range temporal correlations, via detrended fluctuation analysis (DFA) and 5) functional connectivity of EEG data from ES patients (n=40 patients) and healthy controls (n=20 subjects), with multiple blinded measurements during wakefulness and sleep for each patient.

**Results:** In ES patients, EEG amplitude was significantly higher in all electrodes. Shannon and permutation entropy were lower in ES patients than control subjects, while DFA intercept values in ES patients were significantly higher than control subjects. DFA exponent values were not significantly different between the groups. EEG functional connectivity networks in ES patients were significantly stronger than controls. Using logistic regression, a multi-attribute classifier was derived that accurately distinguished cases from controls (area under curve of 0.96).

**Conclusions:** Computational EEG features successfully distinguish ES patients from controls in a large, blinded study.

**Significance:** These objective EEG markers, in combination with other clinical factors, may speed the diagnosis and treatment of the disease, thereby improving long-term outcomes.

**Highlights:**

1. Objective computational EEG features may aid diagnosis of epileptic spasms (ES)
2. ES EEG has increased delta and theta power and decreased entropy relative to controls
3. Stronger functional connectivity networks differentiate ES patients from controls

## I. Introduction

Epileptic spasms (ES) is an epileptic encephalopathy with typical onset between 4-7 months of age (Hrachovy and Frost 2003). These seizures occur in clusters and consist of abrupt muscle spasms, often accompanied by an interictal electroencephalographic (EEG) pattern known as *hypsarrhythmia* (Gibbs et al. 1953; Pavone et al. 2013; Fisher et al. 2018). Hypsarrhythmia is marked by asynchronous, high-amplitude slow waves, a disorganized background, and multi-focal independent spikes (Gibbs et al. 1953; Hrachovy et al. 1981, 1984). Children with ES often exhibit neurocognitive stagnation, psychomotor delay, and other refractory seizure types (Primec et al. 2006; Gaily et al. 2010; Riikonen 2010; Pavone et al. 2013; Widjaja et al. 2015). Prompt, successful treatment increases the likelihood of a favorable outcome, but diagnosis and clinical treatment decisions are challenging and often delayed for various reasons. First, epileptic spasms are associated with a wide range of underlying etiologies (Osborne et al. 2010; Riikonen 2010), and there is potential for misdiagnosis of subtle ES as other seizure types or unrelated benign conditions (Shields 2004; Auvin et al. 2012). Second, the wide variability of EEG patterns, such as hypsarrhythmia and so-called *modified hypsarrhythmia*, can confound visual interpretation of the EEG (Gibbs et al. 1953; Hrachovy et al. 1984; Sue et al. 1997; Frost et al. 2011). Although the presence of hypsarrhythmia is often used as a diagnostic marker of the syndrome, there is low inter-rater reliability for identification of the pattern (Hussain et al. 2015; Mytinger et al. 2015) and it is not a predictor of outcome (Demarest et al. 2017). Overall, rates of sustained treatment response are low due to a paucity of effective first-line therapies and a high relapse rate (Hrachovy et al. 1983; Baram et al. 1996; Ito et al. 2002), and this is further complicated by diagnostic challenges and the subsequent use of inappropriate therapies in cases of misdiagnosis (O’Callaghan et al. 2011; Auvin et al. 2012). Computational EEG biomarkers of ES that are independent of the presence of hypsarrhythmia would help address these challenges by providing tools for objective—and perhaps more accurate— identification of the disease.

We previously demonstrated that several computational metrics are relevant to ES, such as EEG amplitude (Smith et al. 2018), power spectrum (Smith et al. 2018), the strength of long-range temporal correlations in EEG amplitude modulations (Smith et al. 2017), and functional connectivity (Shrey et al. 2018). However, those results were obtained using a relatively small, homogeneous cohort of patients with new-onset ES. Here, we sought to validate these findings in a much larger and more diverse cohort of ES patients, the majority of which exhibited refractory spasms. We measured five types of computational EEG metrics in all subjects: 1) amplitude, 2) power spectrum and spectral edge frequency, 3) Shannon and permutation entropy, 4) long-range temporal correlations, and 5) functional connectivity using both amplitude and phase-based measures. We compared the results to a group of normal children who underwent EEG monitoring to rule out ES or other seizures. In addition to the larger cohort, we have enacted randomization and blinding procedures to mitigate bias in the selection of subjects and EEG clips, and we analyzed EEG in both sleep and wakefulness, with multiple EEG clips for each patient in order to evaluate the reproducibility of computational measurements. This comprehensive study describes objective EEG characteristics that may improve the accuracy and latency of ES diagnosis.

## II. Methods

### 2.1 Identification of Cases and Controls

Approval to perform this study was obtained from the UCLA Institutional Review Board. Using a clinical video-EEG database, which includes all patients who underwent video-EEG monitoring at UCLA Mattel Children’s Hospital between February 2014 and July 2018, we identified ES cases and normal controls as follows. To select cases, we used a computer-aided algorithm to randomly select 50 patients with ES who exhibited epileptic spasms on an overnight video-EEG, regardless of the presence or absence of hypsarrhythmia. To select controls we implemented a similar algorithm to randomly select 25 patients who (1) underwent overnight video-EEG to evaluate for suspected epileptic spasms, (2) exhibited a normal video-EEG, and (3) were deemed neurologically normal (i.e. no known or suspected neurological diagnosis as per clinical neurology assessment after EEG). We chose these criteria to mimic the clinical scenario in which a patient is being evaluated for suspected epileptic spasms, and physicians must determine whether spasms are present or not. Ultimately, we envision our proposed computational EEG features would be used in this situation. However, with these strict inclusion criteria and the high proportion of patients with refractory spasms typically seen at UCLA, we found that the control group was considerably younger than the cases. Therefore, we assembled an approximately age-matched cohort as follows: To add older control patients, we identified 25 children who presented for evaluation of possible seizures (without the criterion that the seizures were epileptic spasms); these patients were ultimately found to be seizure-free and neurologically-normal in a fashion identical to the original control group. We then used an automated algorithm to randomly select subjects (in a 2:1 ratio of cases to controls) from amongst the 50 potential cases and 50 potential controls. To generate similar age distributions in an unbiased fashion, the process was repeated iteratively until the group-wise difference between cases and controls was less than 10% for median age, mean log-transformed age, and standard deviation of log-transformed age. The final cohort included 40 ES cases and 20 control subjects.

### 2.2 Data Abstraction

Two EEG clips during wakefulness and two clips during sleep were extracted for all subjects. Epoch selection was guided by a randomization algorithm such that the abstractor selected EEG data beginning at a specific, predetermined time. Each clip contained 20 minutes of EEG data, and data were sampled at 200 Hz with impedances below 5 kΩ.

### 2.3 EEG artifact identification

Time periods in the EEG containing artifact were identified using an automatic extreme value detection algorithm, similar to previously published methods (Durka et al. 2003; Moretti et al. 2003) (Figure 1). Specifically, the data were first broadband bandpass filtered (1.5 – 40 Hz, Butterworth filter). The mean was subtracted from each channel, and the standard deviation was calculated using the entire zero-mean time series. Whenever the absolute value of the voltage exceeded a threshold of 7.5 standard deviations above the mean value in any single channel, the time points were marked as artifact. A buffer of 0.9 seconds was added before and after the extreme amplitude values to ensure that the entire artifact was marked. Data recorded during EEG impedance checks were also automatically identified and marked. Note that artifacts were identified using broadband filtered data, but the artifactual EEG epochs were removed after the band-specific filtering needed for each metric. In all cases, artifactual segments of data were excluded from all channels, even if the artifact occurred in a single channel.

**Figure 1.**
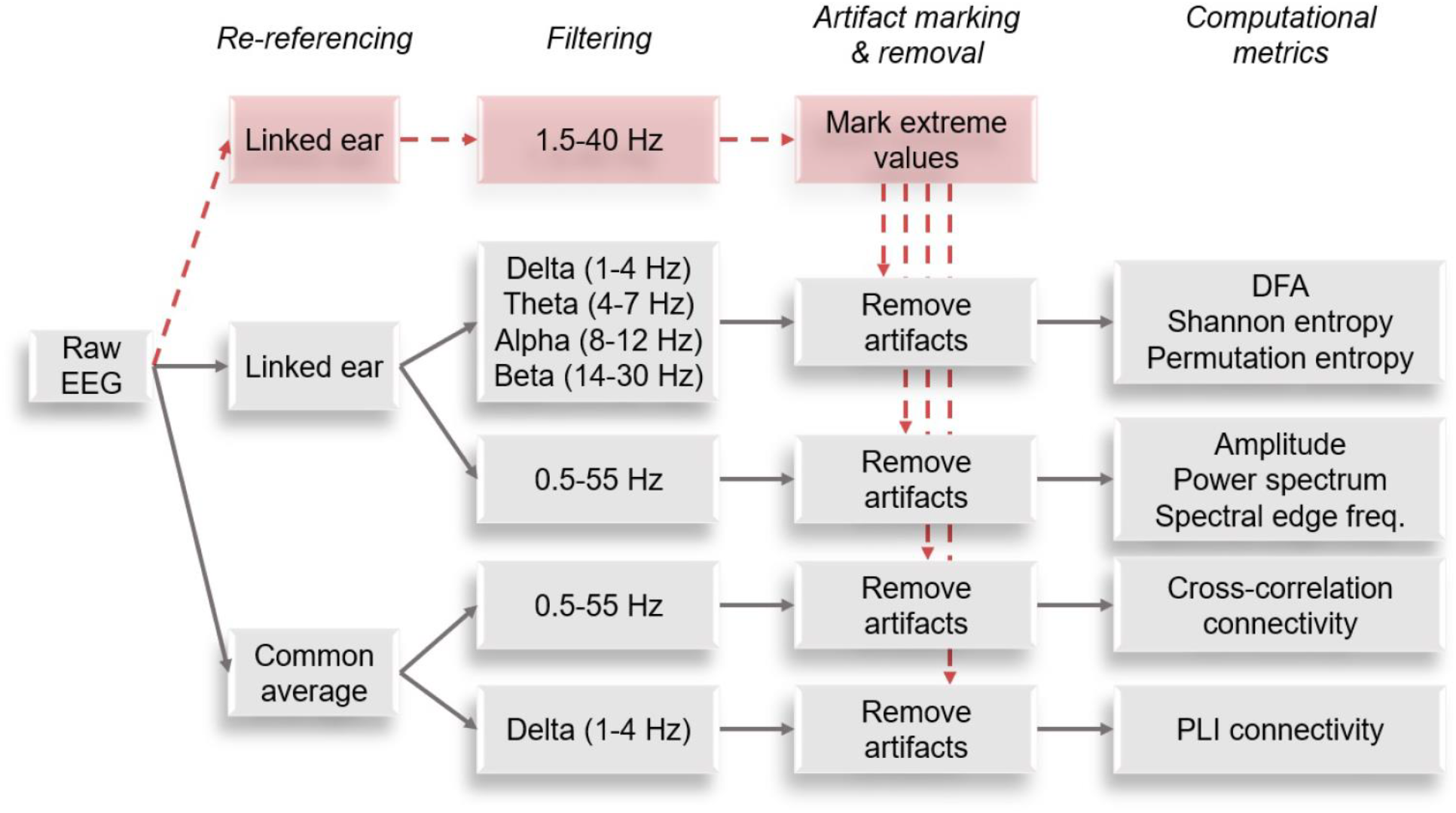
Pre-processing steps to re-reference, filter, and remove artifacts in EEG data from epileptic spasms (ES) patients and control subjects.

### 2.4 Computational EEG metrics

We calculated five types of computational metrics for each EEG recording: amplitude, power spectrum and spectral edge frequency, Shannon and permutation entropy, long-range temporal correlations, and amplitude- and phase-based functional connectivity networks. These metrics were chosen because they are commonly used in EEG signal analysis and demonstrated relevance to ES both in the literature and in our prior studies using a smaller, more homogeneous cohort (Smith et al. 2017, 2018; Shrey et al. 2018). All procedures for data pre-processing, filtering, artifact removal, and calculation of computational metrics are summarized in Figure 1. Data were re-referenced to the common average for the functional connectivity measures, and a linked-ear montage was used for all other analyses (Stam et al. 2007; Shrey et al. 2018). We implemented different filtering strategies as needed for each calculated metric. For calculation of the EEG amplitude, power spectrum, spectral edge frequency, and cross correlation functional connectivity, we bandpass-filtered the re-referenced data from 0.5-55 Hz to include all frequencies of clinical interest. Phase-lag index functional connectivity was measured in the delta frequency band (1-4 Hz); this computational metric requires selection of a narrow frequency range, and the delta band was chosen to enable comparison to the cross-correlation connectivity, which will be primarily driven by the low frequency activity. For calculation of DFA, Shannon entropy, and permutation entropy, we analyzed all standard narrow frequency bands (delta band 1-4 Hz, theta band 4-7 Hz, alpha band 8-12 Hz, and beta band 14-30 Hz).

#### 2.4.1 Amplitude

Because hypsarrhythmia and other interictal patterns are defined by a high amplitude (typically greater than 200-300 μV), the EEG amplitude is a clinically relevant signal feature in ES (Hrachovy et al. 1984; Nehlig et al. 2012; Pavone et al. 2013). Amplitude values were calculated using the range of the broadband filtered data in one second windows for each electrode. The variation of EEG amplitude across channels was visualized via topographic maps. For each group, the topographic maps were constructed using the median value across all patients for each electrode location. The figures were generated using the MATLAB-based EEGLAB function “*topoplot*”. To compare amplitude distributions across subjects, we calculated the empirical cumulative distribution function (CDF) for the Cz electrode in each patient dataset. Electrode Cz was chosen because it is minimally contaminated by artifact.

#### 2.4.2 Power Spectrum and Spectral Edge Frequency

For each channel, data were divided into 5-second epochs, and the power spectrum was calculated via the fast Fourier transform on the broadband bandpass-filtered data (0.5-55 Hz). The mean power spectrum was obtained by averaging the power spectra over all epochs. We calculated the decibel change to compare pathological spectra (in ES subjects) to physiological spectra (control subjects). The dB change is defined as follows:

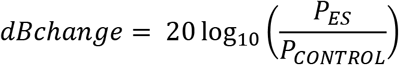

where *P*_*ES*_ is the averaged power spectrum of the ES group and *P*_*CONTROL*_ is the averaged power spectrum of the control group. We report differences in the standard frequency bands of delta (1-4 Hz), theta (4-8 Hz), alpha (8-13 Hz), and beta (13-30 Hz).

To quantify differences in the spectra using a single metric (as opposed to one value for each frequency band), the spectral edge frequency (SEF) was defined as the frequency below which 95% of the power resides (Schwender et al. 1996). Topographic maps of the spectral edge frequency were created by calculating the SEF value for every channel from each patient. Groupwise topographic maps show the median SEF values across subjects, calculated individually for each channel.

#### 2.4.3 Entropy

Entropy is the amount of information contained in a signal, and it is conceptually related to the “predictability” of the data. We calculated both the Shannon entropy and the permutation entropy for ES patients and control subjects.

Shannon entropy is derived from information theory and depends only on the distribution of values in the data; it is independent from the data’s temporal structure. The Shannon entropy H was calculated for each channel as follows (Shannon 1948):

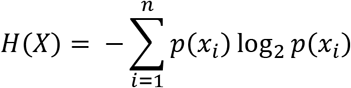

where *p*(*x_i_*) is the probability of observing the *i^th^* value of the bin series in data *x*, and *n* is the number of bins (Cohen 2014). We calculated the optimal number of bins according to Freedman and Diaconis, 1981 (Freedman and Diaconis 1981) as described by Cohen, 2014 (Cohen 2014). The entropy calculation has units of bits, and higher values indicate more stochastic behavior (Van Putten and Stam 2001; Kannathal et al. 2005a, 2005b, 2014; Rosso et al. 2005). We calculated Shannon entropy values for all EEG electrodes and reported the mean entropy value.

Unlike Shannon entropy, permutation entropy quantifies the complexity of a time series while taking the temporal order of the signal into account by the utilization of symbolic dynamics (Bandt and Pompe 2002):

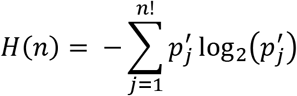

where the 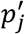 represent the relative frequencies of the possible patterns of symbol sequences and *n* is the embedding dimension (Bandt and Pompe 2002; Riedl et al. 2013). Permutation entropy was calculated on the EEG data from the Cz electrode for each subject in all frequency bands. We used an embedding dimension *n* = 4 and a time delay *τ* = 1.

#### 2.4.4 Long-range temporal correlations

Detrended Fluctuation Analysis (DFA) is a statistical estimation algorithm used to measure the strength of long-range temporal correlations in time series (Peng et al. 1992, 1994; Hardstone et al. 2012). Specifically in neural time series, the temporal modulation of EEG amplitude occurs over periods lasting tens of seconds and is believed to reflect the brain’s ability to control its neuronal synchrony (Linkenkaer-Hansen et al. 2001). Here, DFA was implemented using the algorithm as outlined in our previous study (Smith et al. 2017), adapted from Peng at al. (Peng et al. 1994) and Hardstone et al. (Hardstone et al. 2012). We included box sizes ranging from 1 second to 1/10 of the signal length. If the recording exceeded 1200 seconds in length, the maximum box size was set to 120 seconds. The DFA exponent, denoted *α*, is a direct estimate of the Hurst parameter and reflects the strength of the long-range temporal correlations present in the time series (Hardstone et al. 2012). The *α* value for positively correlated signals varies between 0.5 and 1.0, and human neural electrophysiology data typically falls within this range.

Because we noted little variation across channels, we averaged *α* from all individual channels to obtain a single value for each recording. The intercept of the DFA plot (*β*) was calculated by extrapolating on the logarithmic plot to find the fluctuation value when the window size equaled one sample (the value at which the logarithm of the window size equals zero) (Smith et al. 2017). Similar to the DFA exponent, we averaged *β* from all channels to obtain a single value for each patient. This calculation was done independently for each frequency band.

#### 2.4.5 Functional Connectivity Networks

Functional connectivity is a measure of the correlation between electrophysiological signals in two different brain regions. We calculated functional connectivity networks using both amplitude- and phase-based measures in the ES and control subjects. The amplitude-based measure calculates functional connectivity via cross-correlation in one-second epochs using the method developed by Kramer et al. (Kramer et al. 2009) and Chu et al. (Chu et al. 2012) and previously applied to ES EEG data by our group (Shrey et al. 2018). For each epoch, the connection was deemed to be significant if the maximum cross-correlation value for the channel pair fulfilled two criteria: (1) it occurred at a non-zero lag time, and (2) it exceeded a significance threshold obtained via permutation resampling. The overall connection strength between two channels was calculated as the percentage of epochs in which the connection was significant. For visualization, we created network maps in which edges (connections) between nodes (electrodes) were drawn if the connection value exceeded 0.075. This threshold was not used for any statistical tests.

We additionally applied a phase-based measure of functional connectivity called the phase lag index (PLI). This method calculates the level of synchronization between two electrodes by determining whether the phase of one signal consistently leads or lags the other signal (Stam et al. 2007):

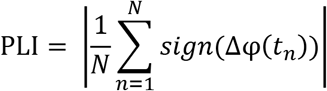

where the PLI value represents the mean signum of the phase difference Δ*φ*(*t*_*n*_) between the two signals over a time period of length N. The instantaneous phase was extracted using the Hilbert transform of the narrow bandpass-filtered EEG signal. We measured PLI between all channels pairs for all 19 electrodes in the delta frequency band (1-4 Hz) in eight-second epochs of clean data, resulting in a 19-by-19 adjacency matrix for each epoch of data. Significance was assessed via surrogate data that was created by shuffling the Fourier phases of the original signal for 100 iterations. The PLI value for the channel pair was considered significant if it exceeded the 95^th^ percentile of the surrogate data distribution. Non-significant connections were replaced with PLI values of zero. The functional connectivity network for each subject was calculated by averaging the pseudo-binary adjacency matrices over all epochs.

### 2.5 Calculation of computational EEG metrics

For each subject, we calculated the metrics for both EEG clips during wakefulness and during sleep, and we averaged the two wakefulness values and the two sleep values. All subsequent analysis was performed with the average wakefulness value and the average sleep value. Note that, for both patient groups, the metric values for the two clips were highly correlated, indicating favorable reproducibility of the measurements (Supplementary Table 1). The metrics were computed on each EEG clip by authors (BAL, DWS, RJS) who were blinded to the patient groups, clip numbers, and designation of wakefulness or sleep.

### 2.6 Statistical Methods

Summary data were reported as mean (standard deviation), or if non-normally distributed, as median (interquartile range). Groupwise comparisons of medians were carried out using the Wilcoxon rank-sum test. Logistic regression models were developed using a forward stepwise approach. Evaluation for multicollinearity was carried out with visual inspection of covariate scatter plots and calculation of all pair-wise correlation coefficients. Adjustment for multiple comparisons was accomplished using the Benjamini-Hochberg procedure. Statistical calculations were carried out using Stata (version 14, Statacorp; College Station, Texas, USA) and MATLAB (ver. 2019b, MathWorks, Inc; Natick, MA, USA).

## III. Results

### 3.1 Subjects

Clinical and demographic attributes of the study population are summarized in Table 1. The median (interquartile range) ages of the 40 cases and 20 controls were 11.0 (7.6 – 22.6) and 9.9 (7.4 – 27.9) months, respectively.

**Table 1.** Clinical and demographic characteristics of the study population

|  |  |  |  |
| --- | --- | --- | --- |
|  | Cases<br>(n = 40) | Controls<br>(n = 20) | p-value<sup>1</sup> |
| Age at EEG, months<sup>2</sup> | 11.0 (7.6, 22.6) | 9.9 (7.4, 27.9) | 0.95 |
| Female, n (%) | 20 (50%) | 8 (40%) | 0.59 |
| Age of onset of epileptic spasms, months<sup>2</sup> | 7.4 (4.6, 11.8) |  |  |
| Duration of epileptic spasms, months<sup>2</sup> | 2.4 (0.1, 6.0) |  |  |
| Normal development at onset of epileptic spasms | 20 (50%) |  |  |
| Known etiology | 31 (78%) |  |  |
| Structural etiology | 22 (55%) |  |  |
| Genetic etiology | 14 (35%) |  |  |

<sup>1</sup>p-values were not adjusted for multiple comparisons.
<sup>2</sup>Median (interquartile range)

### 3.2 High amplitude EEG in ES

During both wakefulness and sleep, the EEG amplitude was higher in ES patients than controls. Significant differences between the distributions of amplitude values can be seen in a representative example from channel Cz (p<0.05, Figure 2A). The differences between ES and control subjects were significant in all channels (BH-adj. p<0.05, Figure 2B), and the spatial variation of the amplitude values is similar between groups (Figure 2B). The highest EEG amplitudes were situated frontally in both groups during wakefulness (Figure 2B, top row), while the highest EEG amplitudes were located more centrally during sleep (Figure 2B, bottom row). Lower amplitudes in the temporal channels may be due to the use of a linked-ear reference.

**Figure 2.**
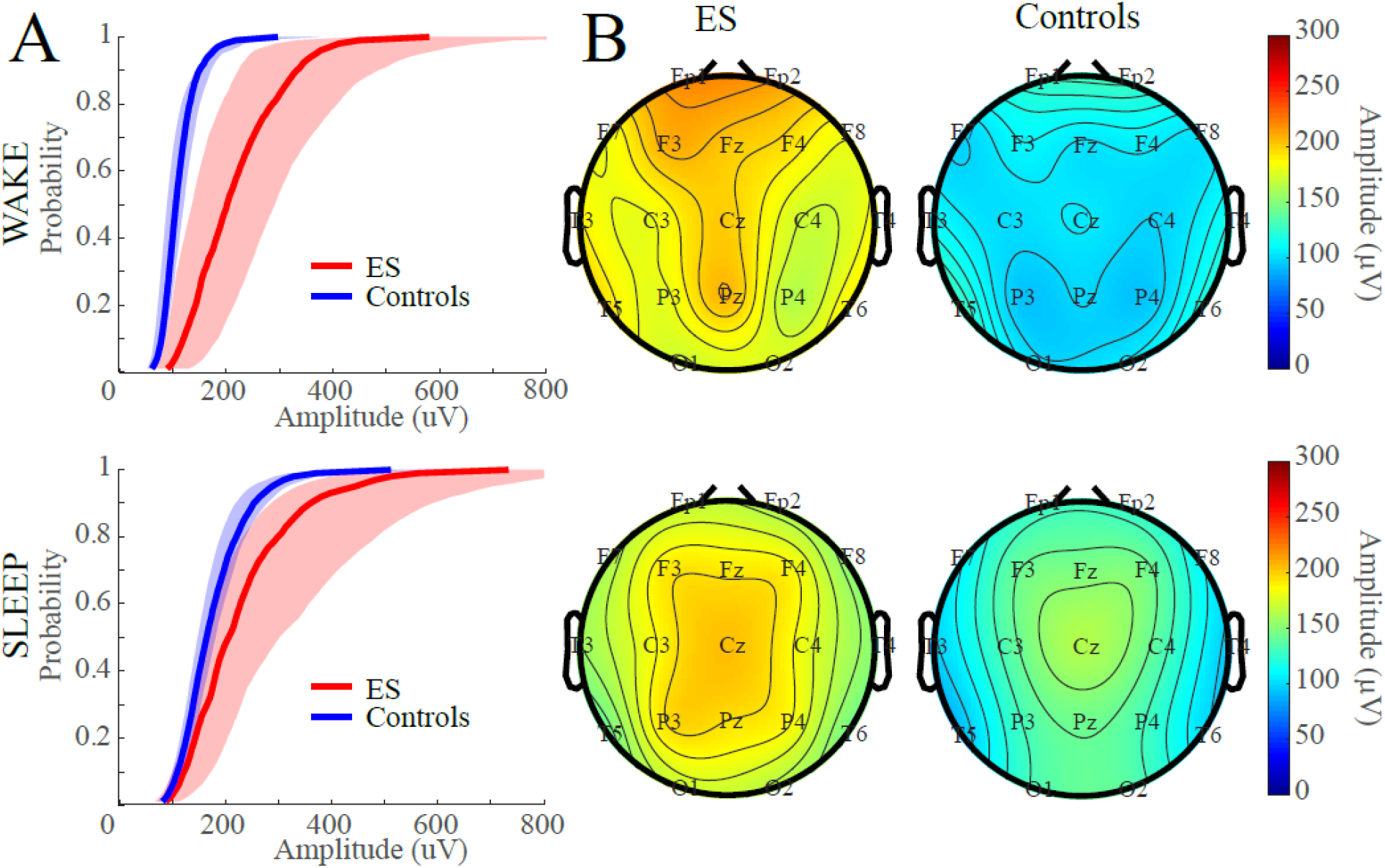
EEG amplitude is higher in epileptic spams (ES) patients than controls. **(A)** Empirical cumulative distribution functions (CDFs) from the Cz electrode during wakefulness (top) and sleep (bottom). The solid line indicates the median of the group CDF values, and the shaded region covers the interquartile range for the ES patients (red) and the controls (blue). **(B)** Topographic maps of median EEG amplitude for ES patients (left column) and controls (right column) during wakefulness (top row) and sleep (bottom row).

### 3.2 High EEG spectral power in ES

During wakefulness and sleep, ES patients exhibited higher power in nearly all frequency bands when compared to the control group (Figure 3A). In wakefulness, the greatest increase in power for ES patients occurs below 10 Hz, while the power in the upper beta frequency band matches that of control subjects (Figure 3A, top). During sleep, ES patient EEG power shows the greatest increase in the lower beta frequency band (14-20 Hz) and the alpha frequency band (8-12 Hz) (Figure 3A, bottom). The raw power spectra for ES patients and control subjects are also provided for reference (Supplementary Figure 1). These differences in the power distribution are reflected by the SEF metric. During wakefulness, the SEF values for ES patients were significantly lower than controls in all channels (BH-adj. p<0.05). The lowest SEF values were in the central head regions in both groups (Figure 3B, top row). During sleep, we found similar spatial variation of SEF values (Figure 3B, bottom row), but the differences were only statistically significant in channels O1 and O2 (BH-adj. p<0.05). Sleep was generally associated with lower SEF values than wakefulness.

**Figure 3.**
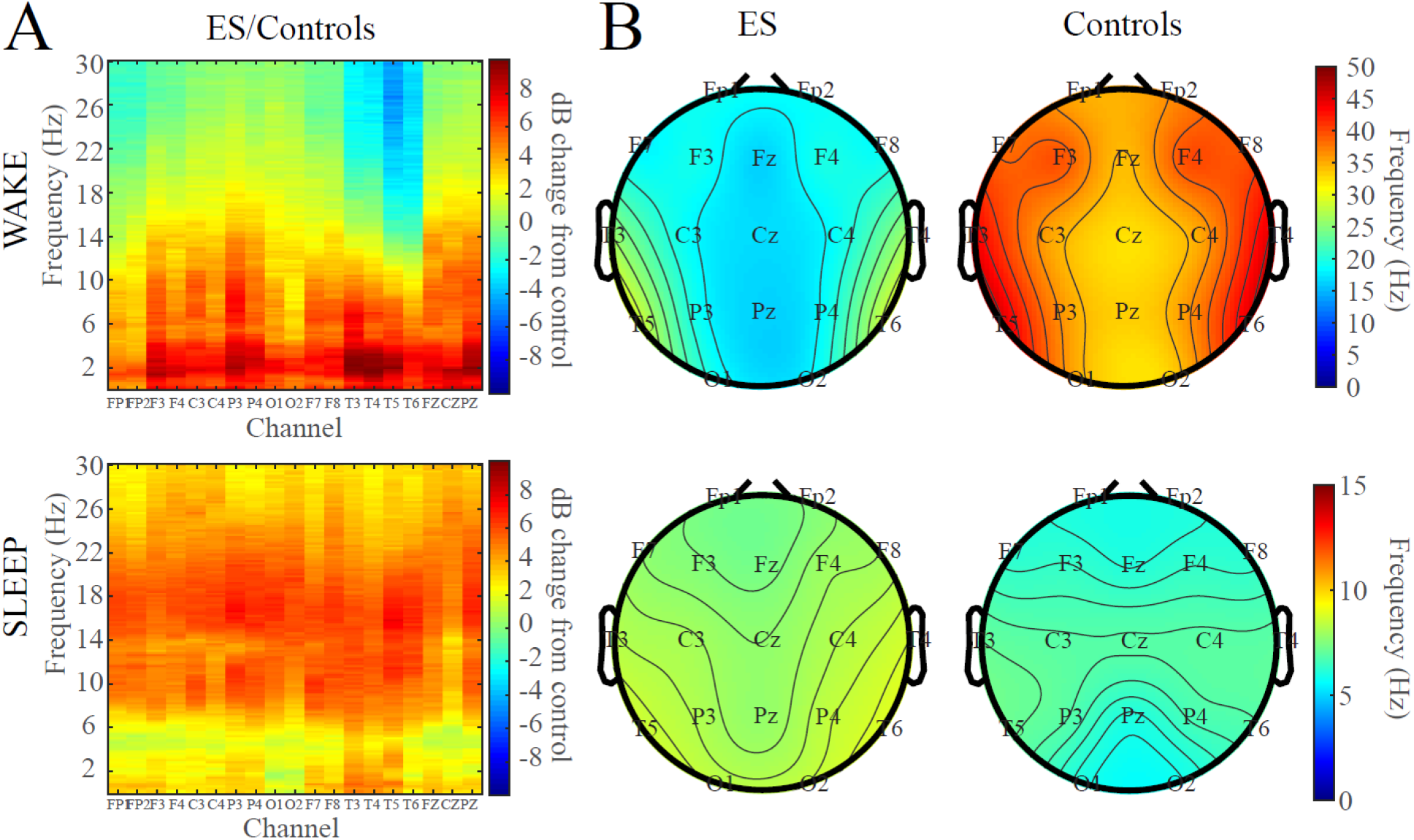
EEG spectral power is higher in epileptic spams (ES) patients than controls. **(A)** EEG power spectra for ES patients, relative to control subjects, during wakefulness (top) and sleep (bottom). **(B)** Topographic maps of spectral edge frequency (SEF) for ES patients (left column) and controls (right column) during wakefulness (top row) and sleep (bottom row).

### 3.3 Low permutation entropy in ES

During wakefulness, Shannon entropy values were significantly higher in ES patients than control subjects in the delta frequency band, but significantly lower in the alpha frequency band (Figure 4A) (BH-adj. p<0.05). Shannon entropy values were also lower for ES patients in all frequency bands during sleep, with the difference being significant in the theta, alpha, and beta frequency bands (BH-adj. p<0.05, Figure 4A).

**Figure 4.**
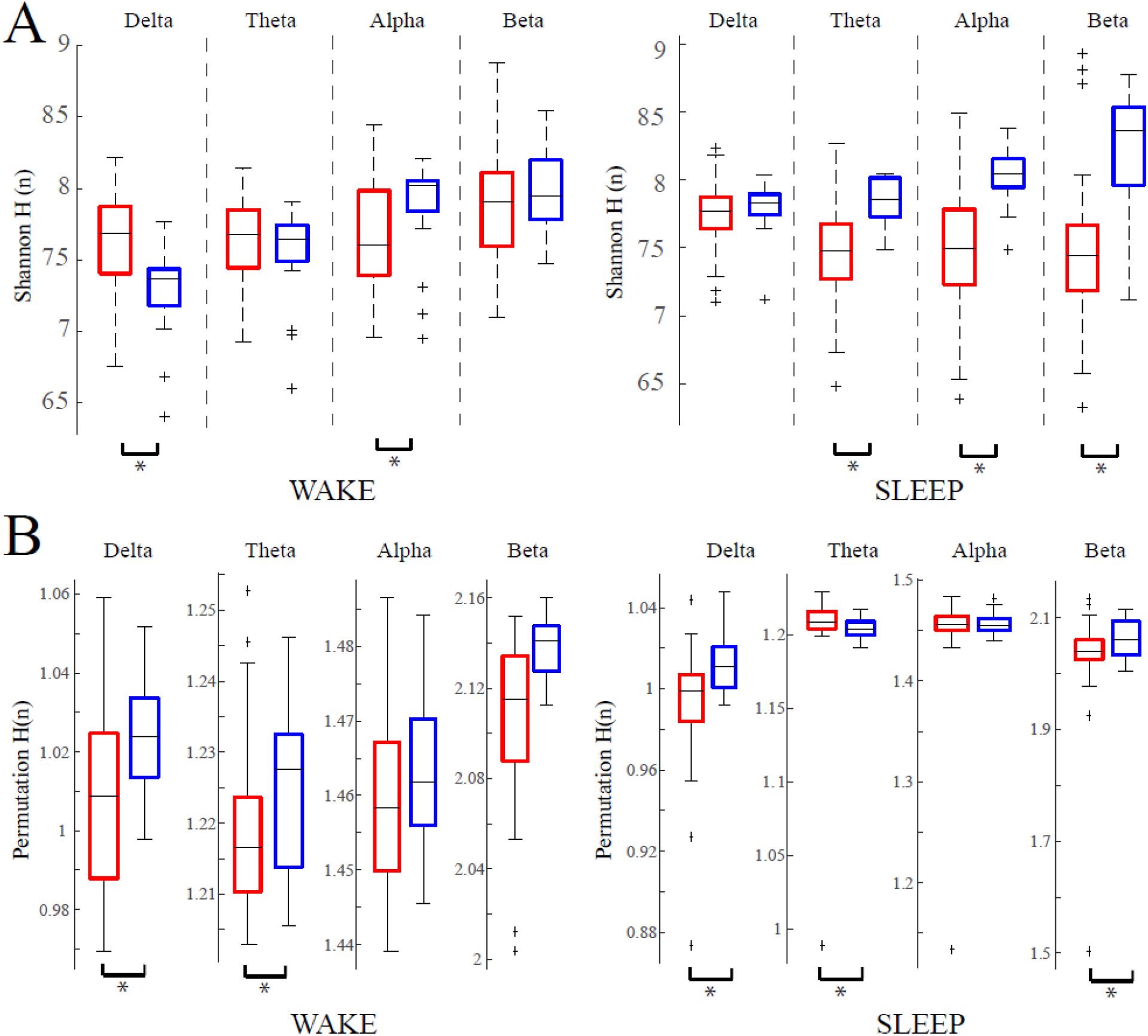
EEG entropy is lower in epileptic spams (ES) patients than controls. **(A)** Shannon entropy and **(B)** permutation entropy for ES patients (red) and control subjects (blue) during wakefulness (left subfigure) and sleep (right subfigure). Each data point represents the mean entropy across channels for a single subject. Asterisks indicate significance of p<0.05.

Similar to the results for Shannon entropy, the permutation entropy was generally lower in ES patients compared to controls. The permutation entropy values were significantly lower in ES patients in the delta and theta frequency bands during wakefulness (Figure 4B) (BH-adj. p<0.05). However, during sleep, the permutation entropy values were significantly lower in the delta and beta frequency bands, but significantly higher in the theta band (Figure 4B) (BH-adj. p<0.05).

### 3.4 Long-range temporal correlations distinguish ES patients from control subjects

To assess differences in the strength of long-range temporal correlations, we plotted the DFA intercept, *β*, against the DFA exponent, *α*, for all groups (Figure 5A). Consistent with our prior work (Smith et al. 2017), we found that *α* and *β* were negatively correlated and that the *β* values for ES patients and controls were largely non-overlapping. To better assess statistical differences between *α* and *β*, we also show boxplots of the DFA exponent (Figure 5B) and DFA intercept (Figure 5C). All differences were tested with a Wilcoxon rank-sum test, corrected via Benjamini-Hochberg procedure with adjusted p<0.05. During wakefulness, the *α* values of each group did not significantly differ from one another in any frequency band (Figure 5B, top row). In sleep, the *α* values for ES patients were significantly higher in the delta band compared to controls (Figure 5B, bottom row). In contrast, the *β* values for ES patients were significantly higher than the control infants in all frequency bands in both awake and sleep data except for the delta frequency band during sleep (Figure 5C).

**Figure 5.**
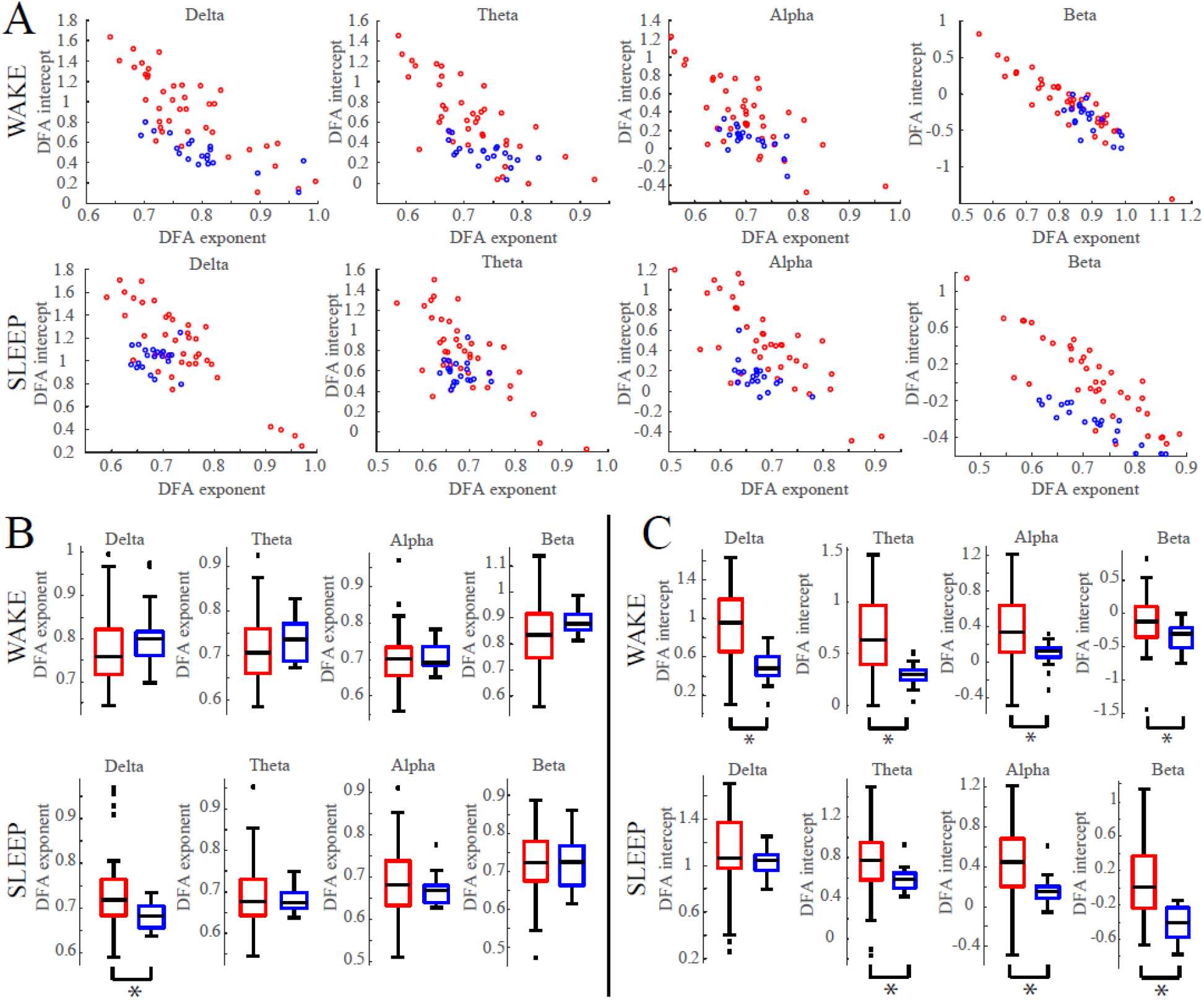
Long-range temporal correlations are altered in epileptic spams (ES) patients; in particular, the detrended fluctuation analysis (DFA) intercept is lower in ES patients compared to controls. **(A)** Scatterplots of DFA parameters in wakefulness (top row) and sleep (bottom row). Points are plotted at (*α*_*i*_, *β*_*i*_) where *α* is the DFA exponent value and *β* is the DFA intercept value for patient *i*. Results are shown for ES (red) and control (blue) subjects in all frequency bands. (B) Same data as in (A), represented as boxplots of DFA exponents and (C) boxplots of DFA intercepts for all frequency bands. Asterisks indicate significance of p<0.05.

### 3.5 Stronger EEG functional connectivity in ES

Overall, we found stronger functional connectivity networks during sleep compared to wakefulness (Figure 6). Using the cross-correlation functional connectivity measure, ES patients exhibited stronger networks compared to controls in both wakefulness and sleep (Figure 6A). During wakefulness, the ES patients exhibited significantly stronger cross-correlation connections than the normal infants in 98 of the 171 possible electrode pairs (57.3%) (Figure 6A, top row) (BH-adj. p<0.05). Similar results were obtained during sleep, with 78 of the 171 possible electrode pairs (45.6%) exhibiting statistically stronger connectivity values than the control group (Figure 6A, bottom row) (BH-adj. p<0.05).

**Figure 6.**
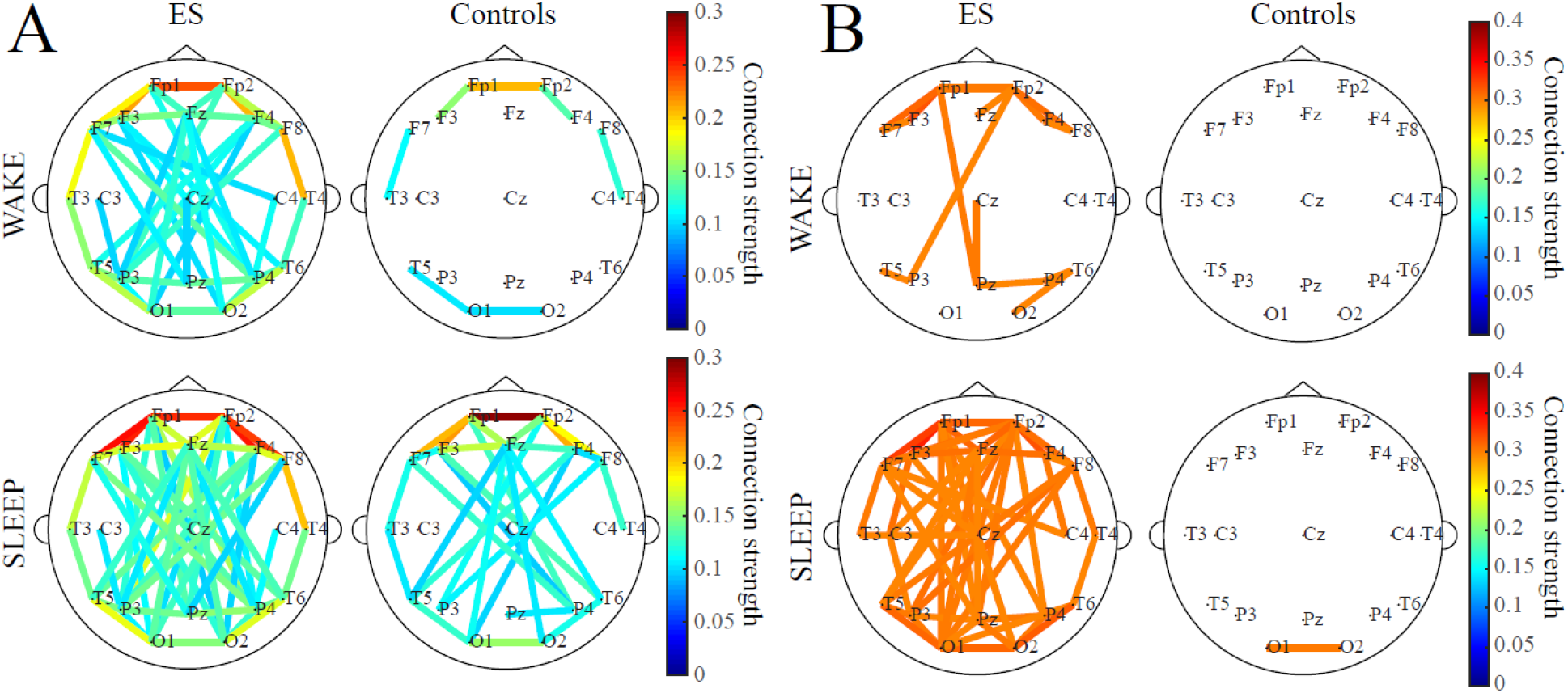
Patients with epileptic spams (ES) have stronger functional connectivity networks than controls. Mean functional connectivity maps are shown for ES patients (left column) and controls (right column) in wakefulness (top row) and sleep (bottom row). **(A)** Cross correlation-based functional connectivity was measured in one-second epochs, and overall connection strength is defined as the proportion of significant one-second epochs. Connection strength is represented by the color of the edges. For visualization, graph edges are displayed if the connection strength between two electrodes exceeds 0.075. **(B)** Functional connectivity networks based on phase-lag index were assessed in eight-second epochs, and statistical significance was tested via phase-shuffled surrogate data. The pseudo-binary matrices were averaged over all available epochs. For visualization, graph edges are displayed if the connectivity strength between two electrodes exceeds 0.2.

The PLI-derived connectivity networks were visually similar in structure to the networks obtained with cross-correlation. Similar to cross-correlation, ES patients exhibited stronger networks when compared to controls in both wakefulness and sleep. In wakefulness, 144/171 (84.2%) of the ES patient functional connections were significantly stronger than the control subjects (Figure 6B, top row) (BH-adj. p<0.05), while 152/171 (88.9%) connections were significantly stronger during sleep (Figure 6B, bottom row) (BH-adj. p<0.05).

### 3.6 Multivariable classification of cases and controls

After establishing group differences in the aforementioned computational EEG metrics, we explored whether a multivariable logistic regression model could be developed to accurately discriminate cases from controls using multiple computational metrics in a simultaneous fashion. Using the age-matched cohorts (40 cases and 20 controls), our model utilized sleep measures of functional connectivity (phase-lag index in the delta frequency band), Shannon entropy in the beta frequency band, and the DFA intercept in the beta band. Using the regression coefficients, we devised a metric, *M*, defined as follows:

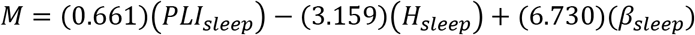

where *PLI* is the phase lag index, *H* is Shannon entropy and *β* is the intercept derived from the DFA analysis. As illustrated in Figure 7 the median *M* was higher among cases than controls (p<0.001), and when evaluated as a classifier with receiver operating characteristic (ROC) analysis, it yielded an area under the curve (AUC) of 96%. In an exploratory internal validation, we evaluated the accuracy with which *M* could discriminate the 10 cases and 30 controls that remained after the selection of the age-matched cohorts. Among these 40 additional patients, median *M* remained higher among cases (0.006), but classification of individual patients was compromised, as suggested by AUC=79%. Recognizing that these remaining cases and controls were age-mismatched and included numerous patients who were far older than typical patients with epileptic spasms, we found that favorable classification was preserved by excluding any of the additional patients whose age was greater than 4 years, as suggested by AUC = 93%. Similarly, considering all 100 candidate cases and controls, while excluding the 4 cases and 22 controls who were older than 4 years, we again observed favorable classification (AUC = 94%). As illustrated in Figure 8, poor classification performance among older patients likely reflects a “pseudomaturation” among cases, such that their entropy and functional connectivity metrics resemble those of older normal controls to some extent.

**Figure 7.**
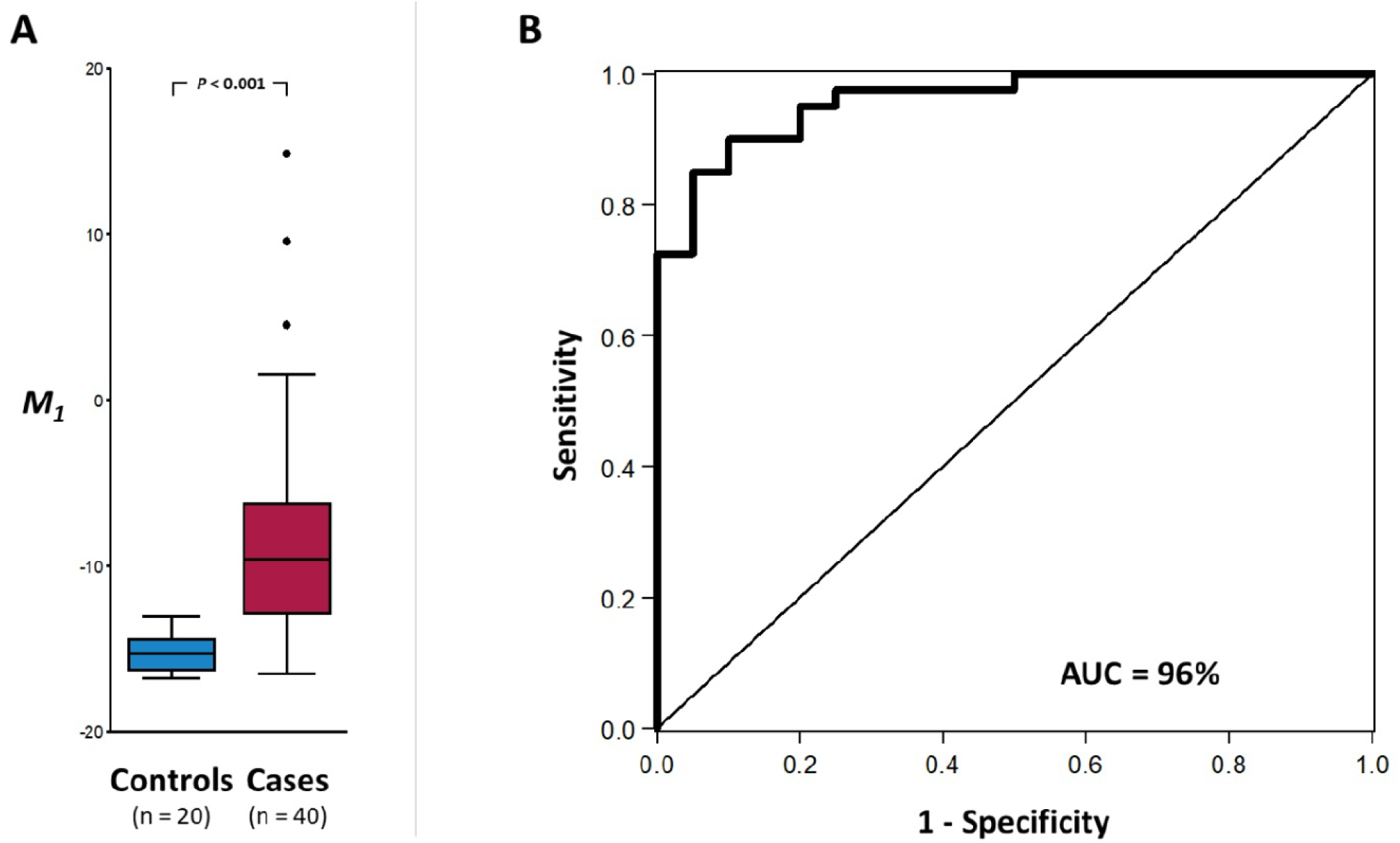
Computational EEG measures enable classification of epileptic spams (ES) patients and controls. **(A)** Values of the multivariable metric, *M*, derived from logistic regression coefficients, for controls (left, blue) and ES patients (right, red). The metric was based on phase-lag index functional connectivity, Shannon entropy, and the detrended fluctuation analysis (DFA) intercept. **(B)** A receiver operating characteristic (ROC) curve was created by sweeping through values of *M* and measuring the accuracy (sensitivity and specificity) of classifying individual subjects as ES patients or controls. The area under the curve was 0.96, indicating that the metric *M* can distinguish between the two groups.

**Figure 8.**
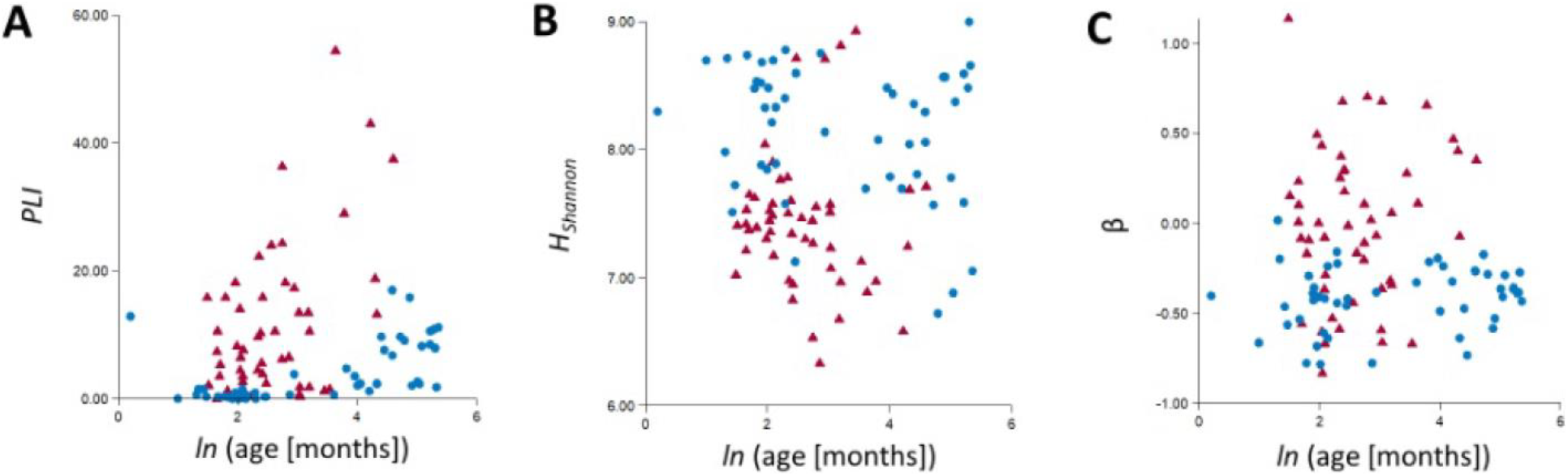
Computational EEG metrics exhibit some dependence on age, particularly for subjects over four years old. The values of **(A)** phase-lag index connectivity, **(B)** Shannon entropy, and **(C)** detrended fluctuation analysis (DFA) intercept, are shown as a function of subject age for epileptic spasms (ES) patients (red triangles) and control subjects (blue circles). Older patients are difficult to classify, as the values of phase-lag index connectivity and Shannon entropy for ES patients overlap with those of control subjects over four years old. Data for all 50 ES patients and 50 control subjects are shown.

## IV. Discussion

We analyzed EEG data from a large, diverse cohort of epileptic spasms patients and identified multiple computational EEG markers that differ significantly from normal infants. This study validates previous work on a smaller, more homogeneous cohort of patients (Smith et al. 2017, 2018; Shrey et al. 2018). The results are more rigorous, as we have included a larger number of subjects, evaluated both sleep and awake EEG recordings, repeated multiple measurements per subject, and used randomization and blinding to mitigate potential bias in the selection of patients and EEG samples. This is the largest and most comprehensive study of its kind.

In our pilot study, we retrospectively identified 21 patients with new-onset ES that were treated at the Children’s Hospital of Orange County and had varying etiologies. The subject age range was narrow (median 6.3, IQR 5.2-8.1 months) and most presented with hypsarrhythmia on the pre-treatment EEG (Smith et al. 2018). In contrast to that study, this cohort included 40 ES patients, with only eight patients exhibiting pre-treatment hypsarrhythmia, and older subjects with a wider age range (median 11.0, IQR 7.6-22.6 months) at the time of treatment. Despite these differences, these two independent studies produced consistent results and suggest that the proposed metrics are robust:

### Amplitude

Amplitude is an EEG feature that is often unusually high in ES (Stamps et al. 1959; Hrachovy et al. 1984; Pavone et al. 2013). Hypsarrhythmia is a high-amplitude pattern, and diffuse slowing is a common feature in both ictal and interictal data (Nehlig et al. 2012). Only eight of the forty patients in this cohort presented with hypsarrhythmia on the pre-treatment EEG; despite that, we found that the EEG had a higher amplitude in ES compared to controls (Table 2), which matches the results from our previous study (Smith et al. 2018). Thus, the differences we report here are not merely reflecting the presence of hypsarrhythmia, suggesting that amplitude may have value as a general biomarker of epileptic spasms.

**Table 2.** Summary of ES results, relative to control subjects, for each computational metric. Abbreviations: Delta frequency band (*δ*), theta frequency band (*θ*), alpha frequency band (*α*), beta frequency band (*β*), N.S. (not significant).

| STATE | *Wake* |  |  |  | *Sleep* |  |  |  |
| --- | --- | --- | --- | --- | --- | --- | --- | --- |
| FREQUENCY BAND | $\delta$ | $\theta$ | $\alpha$ | $\beta$ | $\delta$ | $\theta$ | $\alpha$ | $\beta$ |
| Amplitude | Higher |  |  |  | Higher |  |  |  |
| Power Spectrum | Higher | Higher | Higher | N.S. | Higher | Higher | Higher | Higher |
| Spectral Edge Frequency | Lower |  |  |  | N.S. |  |  |  |
| Shannon Entropy | Higher | N.S. | Lower | N.S. | N.S. | Lower | Lower | Lower |
| Permutation Entropy | Lower | Lower | N.S. | N.S. | Lower | Higher | N.S. | Lower |
| LRTC Strength: Exponent | N.S. | N.S. | N.S. | N.S. | Higher | N.S. | N.S. | N.S. |
| LRTC Strength: Intercept | Higher | Higher | Higher | Higher | N.S. | Higher | Higher | Higher |
| Functional Connectivity: Cross-correlation | Stronger (57.3% of connections) |  |  |  | Stronger (45.6% of connections) |  |  |  |
| Functional Connectivity: Phase-Lag Index | $\delta$ band: Stronger (84.2% of connections) | | | | $\delta$ band: Stronger (88.9% of connections) | | | |

### Power and Spectral Edge Frequency

Consistent with the finding of high amplitude in ES, the EEG power of ES patients was significantly higher than control subjects (Table 2). This is consistent with the clinical findings of diffuse slowing in the pre-treatment EEG (Nehlig et al. 2012). Although we only analyzed frequencies up to 30 Hz in this study, it has been noted that fast activity (14-50 Hz) may play an important role in ES (Inoue et al. 2008; Wu et al. 2008). Even higher frequency ranges (40-150 Hz) may have relevance in ES as well (Kobayashi et al. 2004; Nariai et al. 2020), but the sampling rate of our data precluded this analysis. Particularly during wakefulness, the spectral edge frequency further highlighted the distinction in the frequency characteristics of ES patients in comparison with controls, with lower SEF values signaling significantly higher power in the lower frequency bands. The SEF metric is one way to summarize differences across all frequencies in the power spectra using a single measurement.

### Entropy

Shannon entropy has been reported to be lower in epilepsy patients than healthy subjects (Kannathal et al. 2005a, 2005b, 2014; Rosso et al. 2005). From a nonlinear dynamics perspective, this is because epileptic data often exhibits a lower dimension than healthy data, which is more stochastic in nature. Specifically in ES, it was found that hypsarrhythmia exhibited lower dimension and lower entropy than healthy control data, but the time series was not as nonlinear as seizure data (Van Putten and Stam 2001). Corroborating this literature, we found that the Shannon entropy values during sleep were lower in ES patients when compared to control patients. However, during wakefulness, the Shannon entropy in ES patients was only significantly lower than controls in the alpha frequency band. This may be due to the increased likelihood of residual EEG artifacts during wakefulness. Permutation entropy investigates the organization of the time series while accounting for the short-term temporal structure of the data. Permutation entropy was significantly lower in ES patients compared to control subjects in the delta and theta frequency bands during wakefulness, and in the delta and beta frequency bands during sleep (Table 2).

### Long-range temporal correlations

We used detrended fluctuation analysis (DFA) to compare the temporal structure of the EEG data for ES and control subjects, specifically by measuring power-law scaling and long-range temporal correlations. We previously showed that long-range temporal correlations of EEG amplitude modulations were weaker in new onset ES patients compared to controls, and that DFA exponent values normalized with successful treatment (Smith et al. 2017). In contrast, here we found that the DFA exponents of the ES patients were not significantly different from the control group during wakefulness or sleep, except for the delta frequency band during sleep. The intercept, however, was significantly higher in the ES patient group in comparison to the control group in all frequency bands in wakefulness and in all except the delta band during sleep, generally providing excellent separation between the two groups (Table 2). The DFA intercept scales logarithmically with the variance of the amplitude envelope (Smith et al. 2017), suggesting that the EEG in ES patients not only has a high amplitude, but also that this amplitude varies widely over time.

### Functional connectivity

The EEG patterns in ES have been hypothesized to be driven by neuronal networks including subcortical structures which motivated our analysis of functional connectivity. For example, studies with SPECT (Chiron et al. 1993), PET (Chugani et al. 1992), fMRI (Siniatchkin et al. 2007), and EEG source localization (Japaridze et al. 2013) found that subcortical-cortical interactions may play a role in the development of ES (Chiron et al. 1993). This could explain how diffusely abnormal EEG patterns are observed despite focal etiologies. We assessed functional connectivity in two ways. We first used cross-correlation, which has been shown to reveal stable, patient-specific networks in healthy subjects (Chu et al. 2012) as well as ES patients (Shrey et al. 2018). Previously, long-range, cross-hemispheric connections were observed in ES patients both with coherence (Burroughs et al. 2014) and cross-correlation (Shrey et al. 2018). Here, we found that connectivity strengths were higher in ES in most channel pairs in cross correlation-based networks, and we observed more long-range cross-hemispheric connections in ES compared to controls, corroborating previous work (Table 2) (Burroughs et al. 2014; Shrey et al. 2018). Similarly, using PLI as a second connectivity measure, ES subjects exhibited significantly stronger connections during wakefulness and sleep. We chose to analyze the delta frequency band because it exhibited the greatest power differences between ES patients and control subjects. Moreover, we expected it to correspond most closely to the broadband cross-correlation measure, which had demonstrated relevance in previous studies. Across both subject groups and functional connectivity metrics, the functional connections were stronger during sleep than wakefulness.

### Limitations

Although we attempted to remove as many artifacts as possible, we note that some results may have been affected by residual artifactual data. For example, eye movements and muscle artifacts may have contributed to the high frontal amplitude and higher temporal power observed during wakefulness (Figure 3). Muscle artifact may have contributed to stronger connectivity in the peripheral (hat-band) connections and the skewing of the SEF topological map toward the temporal channels. Additionally, the choice of reference may have affected the metric values; the higher amplitude and lower SEF in the central channels may be due the choice of the linked ear reference. We acknowledge these effects and attempted to minimize their influence on the metric values as much as possible, particularly by comparing to a normal control group that is susceptible to the same types and occurrences of artifacts. Additionally, we note that in some of the metrics, the sleep/wake state played a role in whether the metric successfully distinguished ES patients from control subjects. Further investigation of the differences in the EEG profiles of ES patients during sleep and wakefulness may help identify features that are consistently present regardless of the patient’s sleep/wake state; alternatively, this may suggest to clinicians which specific features are relevant, based on whether the patient is awake or asleep.

### Conclusion

The consistency of the results for ES patients across wide ranges of ages, etiologies, and severity (new onset vs. refractory) indicate that these characteristics of the EEG may be consistently present in epileptic spasms patients. As lead time is one of the strongest prognostic factors in these children, stable metrics that quantify the disease add value to current diagnostic tools. We believe these metrics have the potential to significantly enhance clinical decision making. In particular, as suggested by our logistic-regression derived classifier, these metrics may improve diagnostic accuracy. Future studies will also investigate potential algorithms to identify pre-symptomatic patients, predict response to treatment, measure treatment efficacy, and predict relapse.

## Acknowledgements

This study was sponsored by UCB Biopharma and supported by the CHOC Physician Scientist Scholars Program. UCB Biopharma was not involved in the collection, analysis, and interpretation of the data, or the writing of the manuscript.

## Supplementary Information

### Supplementary Figure

**Supplementary Figure 1.**
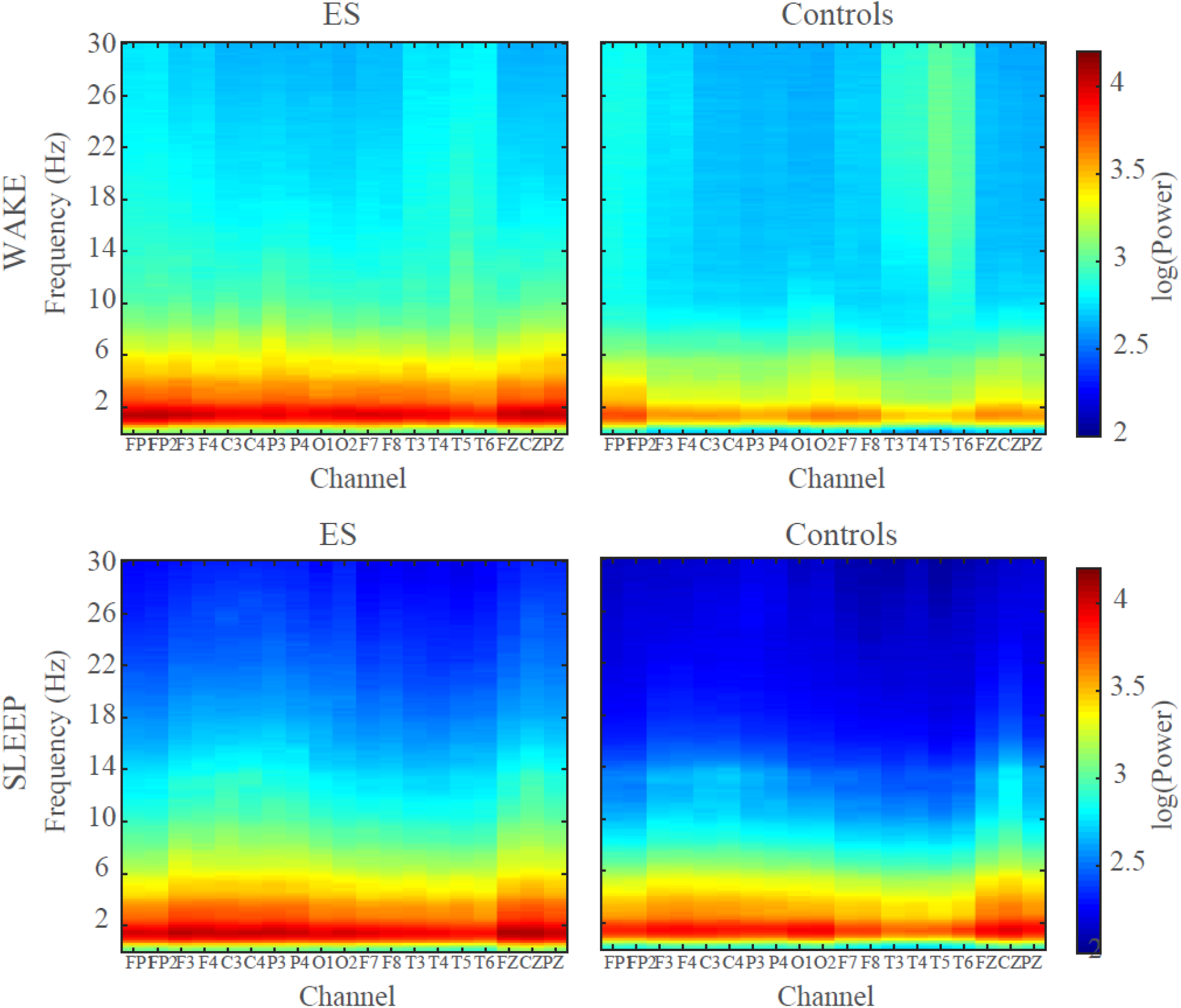
Log-transformed EEG power from 0-30 Hz in all channels for ES patients (left column) and control subjects (right column). EEG power measured during wakefulness is shown on the top row and EEG power measured during sleep is shown on the bottom row.

### Supplementary Tables

**Supplementary Table 1:** Correlation between metrics calculated from independent EEG clips.

| Metric | State | R<sup>2</sup> value | p-value |
| --- | --- | --- | --- |
| DFA exponent: Delta | Wake | 0.51 | 3.9e-296 |
| DFA exponent: Theta | Wake | 0.43 | 1.0e-234 |
| DFA exponent: Alpha | Wake | 0.41 | 1.1e-216 |
| DFA exponent: Beta | Wake | 0.55 | 0 |
| DFA exponent: Delta | Sleep | 0.47 | 1.4e-261 |
| DFA exponent: Theta | Sleep | 0.39 | 1.2e-203 |
| DFA exponent: Alpha | Sleep | 0.42 | 2.6e-228 |
| DFA exponent: Beta | Sleep | 0.46 | 8.2e-258 |
| DFA intercept: Delta | Wake | 0.80 | 0 |
| DFA intercept: Theta | Wake | 0.78 | 0 |
| DFA intercept: Alpha | Wake | 0.74 | 0 |
| DFA intercept: Beta | Wake | 0.59 | 0 |
| DFA intercept: Delta | Sleep | 0.50 | 4.1e-288 |
| DFA intercept: Theta | Sleep | 0.62 | 0 |
| DFA intercept: Alpha | Sleep | 0.64 | 0 |
| DFA intercept: Beta | Sleep | 0.67 | 0 |
| CC Conn | Wake | 0.83 | 3.1e-40 |
| CC Conn | Sleep | 0.81 | 2.7e-38 |
| Amplitude | Wake | 0.63 | 0 |
| Amplitude | Sleep | 0.85 | 0 |
| SEF | Wake | 0.71 | 0 |
| SEF | Sleep | 0.17 | 1.9e-77 |
| Shannon Entropy: Delta | Wake | 0.46 | 7.6e-254 |
| Shannon Entropy: Theta | Wake | 0.33 | 2.8e-169 |
| Shannon Entropy: Alpha | Wake | 0.29 | 8.3e-141 |
| Shannon Entropy: Beta | Wake | 0.28 | 2.0e-137 |
| Shannon Entropy: Delta | Sleep | 0.15 | 3.3e-67 |
| Shannon Entropy: Theta | Sleep | 0.45 | 1.2e-251 |
| Shannon Entropy: Alpha | Sleep | 0.55 | 0 |
| Shannon Entropy: Beta | Sleep | 0.39 | 8.4e-204 |
| Permutation Entropy: Delta | Wake | 0.96 | 6.3e-72 |
| Permutation Entropy: Theta | Wake | 0.99 | 1.2e-101 |
| Permutation Entropy: Alpha | Wake | 0.99 | 7.6e-115 |
| Permutation Entropy: Beta | Wake | 0.98 | 1.4e-93 |
| Permutation Entropy: Delta | Sleep | 0.62 | 2.8e-22 |
| Permutation Entropy: Theta | Sleep | 0.66 | 8.9e-25 |
| Permutation Entropy: Alpha | Sleep | 0.66 | 9.6e-25 |
| Permutation Entropy: Beta | Sleep | 0.65 | 5.4e-24 |
| PLI Conn | Wake | 0.57 | 1.3e-19 |
| PLI Conn | Sleep | 0.51 | 5.9e-17 |

**Supplementary Table 2.** Percentage of EEG clip classified and removed as artifact.

| Percentage Artifact | Wake 1 | Sleep 1 | Wake 2 | Sleep 2 |
| --- | --- | --- | --- | --- |
| ES Patient 1 | 10.92 | 0.22 | 10.84 | 0.50 |
| ES Patient 2 | 23.00 | 5.43 | 22.84 | 3.37 |
| ES Patient 3 | 8.12 | 1.20 | 8.76 | 2.07 |
| ES Patient 4 | 5.69 | 4.06 | 5.37 | 6.25 |
| ES Patient 5 | 13.62 | 1.04 | 11.54 | 2.08 |
| ES Patient 6 | 1.80 | 0.00 | 2.62 | 0.00 |
| ES Patient 7 | 10.49 | 3.41 | 22.00 | 5.88 |
| ES Patient 8 | 11.40 | 1.84 | 19.15 | 2.35 |
| ES Patient 9 | 5.32 | 0.15 | 4.14 | 0.35 |
| ES Patient 10 | 9.03 | 0.66 | 8.17 | 0.20 |
| ES Patient 11 | 13.02 | 1.52 | 10.60 | 1.25 |
| ES Patient 12 | 4.63 | 1.30 | 10.05 | 2.70 |
| ES Patient 13 | 5.23 | 0.06 | 4.23 | 0.00 |
| ES Patient 14 | 12.38 | 1.23 | 9.81 | 0.10 |
| ES Patient 15 | 18.68 | 2.47 | 19.71 | 2.60 |
| ES Patient 16 | 5.72 | 6.08 | 0.10 | 0.00 |
| ES Patient 17 | 8.89 | 0.30 | 8.88 | 0.55 |
| ES Patient 18 | 4.43 | 4.94 | 9.16 | 3.03 |
| ES Patient 19 | 7.98 | 12.79 | 10.84 | 13.19 |
| ES Patient 20 | 2.61 | 0.12 | 5.14 | 0.25 |
| ES Patient 21 | 0.00 | 0.21 | 2.05 | 0.24 |
| ES Patient 22 | 4.83 | 3.52 | 13.12 | 1.93 |
| ES Patient 23 | 14.13 | 0.25 | 10.42 | 0.56 |
| ES Patient 24 | 11.04 | 1.28 | 13.53 | 0.26 |
| ES Patient 25 | 0.35 | 3.62 | 2.49 | 2.00 |
| ES Patient 26 | 13.37 | 0.96 | 14.67 | 0.90 |
| ES Patient 27 | 15.42 | 0.50 | 17.94 | 4.09 |
| ES Patient 28 | 19.50 | 0.91 | 12.71 | 1.45 |
| ES Patient 29 | 5.33 | 3.03 | 15.97 | 2.64 |
| ES Patient 30 | 5.52 | 0.24 | 1.70 | 0.93 |
| ES Patient 31 | 3.25 | 9.82 | 8.13 | 5.82 |
| ES Patient 32 | 8.15 | 5.06 | 14.44 | 4.48 |
| ES Patient 33 | 11.39 | 3.50 | 9.22 | 4.91 |
| ES Patient 34 | 11.25 | 7.86 | 7.39 | 4.39 |
| ES Patient 35 | 16.22 | 4.58 | 18.26 | 0.00 |
| ES Patient 36 | 8.36 | 2.02 | 12.98 | 2.05 |
| ES Patient 37 | 1.59 | 0.21 | 0.90 | 0.10 |
| ES Patient 38 | 11.63 | 2.81 | 9.68 | 0.72 |
| ES Patient 39 | 2.43 | 2.27 | 0.82 | 2.71 |
| ES Patient 40 | 11.62 | 0.44 | 16.10 | 1.24 |
| ES Patient 41 | 2.16 | 0.20 | 0.66 | 0.60 |
| ES Patient 42 | 5.19 | 6.55 | 4.68 | 4.08 |
| ES Patient 43 | 3.51 | 0.83 | 5.21 | 4.09 |
| ES Patient 44 | 13.26 | 1.93 | 13.88 | 4.05 |
| ES Patient 45 | 2.69 | 0.90 | 1.19 | 1.78 |
| ES Patient 46 | 10.55 | 10.05 | 6.82 | 6.28 |
| ES Patient 47 | 1.68 | 0.15 | 0.42 | 0.00 |
| ES Patient 48 | 2.81 | 2.07 | 3.98 | 2.41 |
| ES Patient 49 | 6.70 | 5.12 | 1.83 | 4.61 |
| ES Patient 50 | 5.65 | 1.39 | 5.56 | 2.38 |
| Control 1 | 5.66 | 1.72 | 11.78 | 0.16 |
| Control 2 | 31.18 | 5.53 | 17.44 | 2.36 |
| Control 3 | 8.33 | 3.22 | 9.47 | 0.50 |
| Control 4 | 14.96 | 0.05 | 16.29 | 0.00 |
| Control 5 | 11.52 | 0.53 | 13.27 | 0.48 |
| Control 6 | 10.50 | 1.69 | 8.58 | 0.17 |
| Control 7 | 6.86 | 0.33 | 10.98 | 0.00 |
| Control 8 | 7.32 | 1.02 | 6.68 | 0.31 |
| Control 9 | 12.06 | 0.58 | 18.61 | 0.48 |
| Control 10 | 5.10 | 0.63 | 3.54 | 0.52 |
| Control 11 | 12.35 | 0.31 | 2.93 | 1.87 |
| Control 12 | 18.01 | 0.56 | 8.15 | 2.39 |
| Control 13 | 4.62 | 0.15 | 3.36 | 0.15 |
| Control 14 | 7.14 | 0.14 | 9.71 | 1.11 |
| Control 15 | 2.77 | 0.00 | 4.33 | 0.30 |
| Control 16 | 15.04 | 0.91 | 9.47 | 0.00 |
| Control 17 | 6.61 | 1.51 | 10.02 | 1.88 |
| Control 18 | 8.26 | 0.17 | 12.13 | 0.15 |
| Control 19 | 8.96 | 0.25 | 8.09 | 0.00 |
| Control 20 | 13.34 | 0.71 | 13.15 | 0.80 |
| Control 21 | 4.64 | 0.54 | 3.74 | 0.90 |
| Control 22 | 6.38 | 1.52 | 7.81 | 2.39 |
| Control 23 | 8.61 | 0.00 | 13.30 | 0.00 |
| Control 24 | 20.83 | 0.71 | 6.62 | 0.00 |
| Control 25 | 29.46 | 0.80 | 12.26 | 0.43 |
| Control 26 | 30.59 | 0.52 | 22.52 | 0.00 |
| Control 27 | 4.26 | 0.00 | 1.98 | 0.00 |
| Control 28 | 26.80 | 0.37 | 33.50 | 0.74 |
| Control 29 | 25.74 | 0.00 | 7.27 | 0.10 |
| Control 30 | 5.67 | 2.55 | 7.07 | 1.43 |
| Control 31 | 9.86 | 0.91 | 6.45 | 0.57 |
| Control 32 | 21.79 | 0.53 | 13.57 | 1.55 |
| Control 33 | 14.80 | 0.40 | 7.31 | 0.70 |
| Control 34 | 9.42 | 0.70 | 7.20 | 1.13 |
| Control 35 | 5.91 | 0.15 | 15.72 | 1.11 |
| Control 36 | 9.85 | 1.15 | 7.88 | 0.54 |
| Control 37 | 7.10 | 0.00 | 4.91 | 0.37 |
| Control 38 | 8.56 | 0.10 | 6.61 | 0.31 |
| Control 39 | 9.89 | 0.10 | 8.67 | 0.20 |
| Control 40 | 11.53 | 0.90 | 5.11 | 0.29 |
| Control 41 | 8.66 | 0.98 | 9.67 | 3.56 |
| Control 42 | 19.80 | 2.39 | 23.26 | 1.06 |
| Control 43 | 9.44 | 0.61 | 8.72 | 0.94 |
| Control 44 | 7.30 | 0.00 | 9.66 | 0.65 |
| Control 45 | 15.06 | 0.00 | 7.84 | 0.63 |
| Control 46 | 8.52 | 0.00 | 5.02 | 0.57 |
| Control 47 | 18.66 | 1.86 | 19.71 | 2.32 |
| Control 48 | 4.43 | 0.75 | 4.35 | 0.97 |
| Control 49 | 9.03 | 0.33 | 4.04 | 0.75 |
| Control 50 | 12.45 | 1.67 | 11.72 | 0.00 |

**Supplementary Table 3:** Number of 5-second windows used in power spectrum analysis.

| **Number of 5 second windows used in power spectrum analysis** | **Wake 1** | **Sleep 1** | **Wake 2** | **Sleep 2** |
| --- | --- | --- | --- | --- |
| ES Patient 1 | 322 | 332 | 302 | 359 |
| ES Patient 2 | 239 | 330 | 277 | 342 |
| ES Patient 3 | 354 | 360 | 331 | 349 |
| ES Patient 4 | 343 | 344 | 335 | 332 |
| ES Patient 5 | 233 | 353 | 311 | 266 |
| ES Patient 6 | 371 | 318 | 271 | 362 |
| ES Patient 7 | 323 | 241 | 334 | 230 |
| ES Patient 8 | 318 | 347 | 299 | 356 |
| ES Patient 9 | 199 | 235 | 83 | 314 |
| ES Patient 10 | 355 | 316 | 280 | 366 |
| ES Patient 11 | 176 | 356 | 311 | 349 |
| ES Patient 12 | 347 | 424 | 318 | 347 |
| ES Patient 13 | 343 | 356 | 344 | 360 |
| ES Patient 14 | 317 | 354 | 323 | 367 |
| ES Patient 15 | 152 | 350 | 290 | 344 |
| ES Patient 16 | 338 | 200 | 357 | 349 |
| ES Patient 17 | 286 | 240 | 327 | 329 |
| ES Patient 18 | 199 | 278 | 153 | 301 |
| ES Patient 19 | 330 | 267 | 276 | 235 |
| ES Patient 20 | 99 | 310 | 223 | 288 |
| ES Patient 21 | 87 | 197 | 244 | 327 |
| ES Patient 22 | 114 | 266 | 172 | 349 |
| ES Patient 23 | 308 | 363 | 301 | 362 |
| ES Patient 24 | 188 | 224 | 203 | 280 |
| ES Patient 25 | 104 | 126 | 224 | 358 |
| ES Patient 26 | 158 | 353 | 209 | 352 |
| ES Patient 27 | 214 | 325 | 241 | 295 |
| ES Patient 28 | 257 | 359 | 317 | 351 |
| ES Patient 29 | 220 | 227 | 170 | 228 |
| ES Patient 30 | 141 | 296 | 189 | 348 |
| ES Patient 31 | 305 | 317 | 317 | 344 |
| ES Patient 32 | 158 | 346 | 159 | 269 |
| ES Patient 33 | 127 | 237 | 321 | 221 |
| ES Patient 34 | 228 | 334 | 328 | 313 |
| ES Patient 35 | 296 | 224 | 309 | 201 |
| ES Patient 36 | 213 | 331 | 298 | 349 |
| ES Patient 37 | 197 | 354 | 351 | 352 |
| ES Patient 38 | 315 | 148 | 287 | 202 |
| ES Patient 39 | 244 | 163 | 231 | 209 |
| ES Patient 40 | 277 | 329 | 307 | 387 |
| ES Patient 41 | 186 | 355 | 163 | 303 |
| ES Patient 42 | 326 | 337 | 354 | 242 |
| ES Patient 43 | 156 | 217 | 222 | 191 |
| ES Patient 44 | 204 | 169 | 207 | 228 |
| ES Patient 45 | 185 | 241 | 93 | 202 |
| ES Patient 46 | 138 | 157 | 226 | 162 |
| ES Patient 47 | 177 | 238 | 269 | 249 |
| ES Patient 48 | 143 | 174 | 214 | 227 |
| ES Patient 49 | 139 | 174 | 176 | 109 |
| ES Patient 50 | 155 | 185 | 177 | 141 |
| Control 1 | 252 | 229 | 219 | 231 |
| Control 2 | 161 | 272 | 202 | 251 |
| Control 3 | 218 | 228 | 219 | 240 |
| Control 4 | 300 | 374 | 304 | 354 |
| Control 5 | 263 | 309 | 313 | 229 |
| Control 6 | 214 | 190 | 238 | 211 |
| Control 7 | 223 | 222 | 245 | 240 |
| Control 8 | 219 | 164 | 223 | 241 |
| Control 9 | 221 | 252 | 197 | 234 |
| Control 10 | 206 | 242 | 238 | 223 |
| Control 11 | 275 | 353 | 3485 | 334 |
| Control 12 | 188 | 252 | 226 | 263 |
| Control 13 | 347 | 322 | 334 | 326 |
| Control 14 | 224 | 253 | 214 | 247 |
| Control 15 | 249 | 281 | 264 | 243 |
| Control 16 | 305 | 357 | 335 | 329 |
| Control 17 | 235 | 237 | 225 | 246 |
| Control 18 | 219 | 214 | 211 | 245 |
| Control 19 | 205 | 242 | 224 | 240 |
| Control 20 | 201 | 204 | 218 | 172 |
| Control 21 | 232 | 235 | 227 | 207 |
| Control 22 | 241 | 248 | 224 | 309 |
| Control 23 | 274 | 275 | 240 | 238 |
| Control 24 | 278 | 254 | 335 | 348 |
| Control 25 | 248 | 344 | 296 | 336 |
| Control 26 | 244 | 323 | 262 | 350 |
| Control 27 | 354 | 354 | 366 | 368 |
| Control 28 | 266 | 362 | 242 | 245 |
| Control 29 | 217 | 350 | 309 | 356 |
| Control 30 | 337 | 347 | 334 | 354 |
| Control 31 | 317 | 237 | 324 | 353 |
| Control 32 | 283 | 363 | 324 | 351 |
| Control 33 | 305 | 474 | 330 | 362 |
| Control 34 | 323 | 351 | 324 | 346 |
| Control 35 | 223 | 265 | 198 | 237 |
| Control 36 | 318 | 350 | 327 | 337 |
| Control 37 | 303 | 350 | 335 | 297 |
| Control 38 | 271 | 349 | 331 | 348 |
| Control 39 | 271 | 354 | 331 | 367 |
| Control 40 | 315 | 282 | 359 | 376 |
| Control 41 | 325 | 273 | 289 | 346 |
| Control 42 | 284 | 348 | 262 | 376 |
| Control 43 | 313 | 328 | 332 | 350 |
| Control 44 | 315 | 337 | 322 | 319 |
| Control 45 | 305 | 362 | 321 | 347 |
| Control 46 | 325 | 334 | 341 | 256 |
| Control 47 | 279 | 227 | 266 | 343 |
| Control 48 | 340 | 339 | 334 | 322 |
| Control 49 | 199 | 222 | 241 | 237 |
| Control 50 | 220 | 226 | 213 | 260 |

## REFERENCES

Auvin S, Hartman AL, Desnous B, Moreau AC, Alberti C, Delanoe C, et al. Diagnosis delay in West syndrome: misdiagnosis and consequences. Eur J Pediatr. 2012;171(11):1695–701.

Bandt C, Pompe B. Permutation entropy: a natural complexity measure for time series. Phys Rev Lett. 2002;88(17):174102.

Baram TZ, Mitchell WG, Tournay A, Snead III OC, Hanson RA, Horton EJ. High-dose Corticotropin (ACTH) verses Prednisone for Infantile Spasms: A Prospective, Randomized, Blinded Study. Pediatrics. 1996;97(3):375–9.

Burroughs SA, Morse RP, Mott SH, Holmes GL. Brain connectivity in West syndrome. Seizure Eur J Epilepsy [Internet]. 2014;23(7):576–9. Available from: http://dx.doi.org/10.1016/j.seizure.2014.03.016

Chiron C, Dulac O, Bulteau C, Nuttin C, Depas G, Raynaud C, et al. Study of Regional Cerebral Blood Flow in West Syndrome. Epilepsia. 1993;34(4):707–15.

Chu CJ, Kramer MA, Pathmanathan J, Bianchi MT, Westover MB, Wizon L, et al. Emergence of Stable Functional Networks in Long-Term Human Electroencephalography. J Neurosci. 2012;32(8):2703–13.

Chugani HT, Shewmon DA, Sankar R, Chen BC, Phelps ME. Infantile spasms: II. Lenticular nuceli and brain stem activation on positron emission tomography. Ann Neurol. 1992;31(2):212–9.

Cohen MX. Analyzing Neural Time Series Data: Theory and Practice. Cambridge, Massachusetts: The MIT Press; 2014.

Demarest ST, Shellhaas RA, Gaillard WD, Keator C, Nickels KC, Hussain SA, et al. The impact of hypsarrhythmia on infantile spasms treatment response: Observational cohort study from the National Infantile Spasms Consortium. Epilepsia. 2017;58(12):2098–103.

Durka PJ, Klekowicz H, Blinowska KJ, Szelenberger W, Niemcewicz S. A simple system for detection of EEG artifacts in polysomnographic recordings. IEEE Trans Biomed Eng. 2003;50(4):526–8.

Fisher RS, Cross JH, French JA, Higurashi N, Hirsch E, Jansen FE, et al. Operational classification of seizure types by the International League Against Epilepsy: position paper of the ILAE Commission for Classification and Terminology. Zeitschrift fur Epileptol. 2018;31(4):272–81.

Freedman D, Diaconis P. On the histogram as a density estimator: L2 theory. Probab Theory Relat Fields. 1981;57(4):453–76.

Frost JD, Lee CL, Hrachovy RA, Swann JW. High frequency EEG activity associated with ictal events in an animal model of infantile spasms. Epilepsia. 2011;52(1):53–62.

Gaily E, Liukkonen E, Paetau R, Rekola M, Granström M-L. Infantile spasms: diagnosis and assessment of treatment response by video-EEG. Dev Med Child Neurol. 2010;43(10):658–67.

Gibbs EL, Fleming MM, Gibbs FA. Diagnosis and Prognosis of Hypsarrhythmia and Infantile Spasms. Pediatrics. 1953;13(1):66–73.

Hardstone R, Poil SS, Schiavone G, Jansen R, Nikulin V V., Mansvelder HD, et al. Detrended fluctuation analysis: A scale-free view on neuronal oscillations. Front Physiol. 2012;3(450):75–87.

Hrachovy RA, Frost JD. Infantile Epileptic Encephalopathy with Hypsarrhythmia (Infantile spasms/West syndrome). J Clin Neurophysiol. 2003;20(6):408–25.

Hrachovy RA, Frost JD, Kellaway P. Sleep characteristics in infantile spasms. Neurology. 1981;31(6).

Hrachovy RA, Frost JD, Kellaway P. Hypsarrhythmia: variations on the theme. Epilepsia. 1984;25(3):317–25.

Hrachovy RA, Frost JD, Kellaway P, Zion TE. Double-blind study of ACTH vs prednisone therapy in infantile spasms. J Pediatr. 1983;103(4):641–5.

Hussain SA, Kwong G, Millichap JJ, Mytinger JR, Ryan N, Matsumoto JH, et al. Hypsarrhythmia assessment exhibits poor interrater reliability: A threat to clinical trial validity. Epilepsia. 2015;56(1):77–81.

Inoue T, Kobayashi K, Oka M, Yoshinaga H, Ohtsuka Y. Spectral characteristics of EEG gamma rhythms associated with epileptic spasms. Brain Dev. 2008;30(5):321–8.

Ito M, Aiba H, Hashimoto K, Kuroki S, Tomiwa K, Okuno T. Low-dose ACTH therapy for West syndrome: Initial effects and long-term outcome. Neurology. 2002;58:110–4.

Japaridze N, Muthuraman M, Moeller F, Boor R, Anwar AR, Deuschl G, et al. Neuronal networks in west syndrome as revealed by source analysis and renormalized partial directed coherence. Brain Topogr. 2013;26(1):157–70.

Kannathal N, Acharya UR, Lim CM, Sadasivan PK. Characterization of EEG - A comparative study. Comput Methods Programs Biomed. 2005a;80(1):17–23.

Kannathal N, Chee J, Er K, Lim K, Tat OH. Chaotic Analysis of Epileptic EEG Signals. In: Goh J, editor. The 15th International Conference on Biomedical Engineering. Geneva: Springer International Publishing; 2014. p. 652–4.

Kannathal N, Choo ML, Acharya UR, Sadasivan PK. Entropies for detection of epilepsy in EEG. Comput Methods Programs Biomed. 2005b;80(3):187–94.

Kobayashi K, Oka M, Akiyama T, Inoue T, Abiru K, Ogino T, et al. Very Fast Rhythmic Activity on Scalp EEG Associated with Epileptic Spasms. Epilepsia. 2004;45(5):488–96.

Kramer MA, Eden UT, Cash SS, Kolaczyk ED. Network interference with confidence from multivariate time series. Phys Rev E [Internet]. 2009;79(6). Available from: http://journals.aps.org/pre/pdf/10.1103/PhysRevE.79.061916

Linkenkaer-Hansen K, Nikouline V V, Palva JM, Ilmoniemi RJ. Long-range temporal correlations and scaling behavior in human brain oscillations. J Neurosci. 2001;21(4):1370–7.

Moretti D V., Babiloni F, Carducci F, Cincotti F, Remondini E, Rossini PM, et al. Computerized processing of EEG-EOG-EMG artifacts for multi-centric studies in EEG oscillations and event-related potentials. Int J Psychophysiol. 2003;47(3):199–216.

Mytinger JR, Hussain SA, Islam MP, Millichap JJ, Patel AD, Ryan NR, et al. Improving the inter-rater agreement of hypsarrhythmia using a simplified EEG grading scale for children with infantile spasms. Epilepsy Res [Internet]. 2015;116:93–8. Available from: http://dx.doi.org/10.1016/j.eplepsyres.2015.07.008

Nariai H, Hussain SA, Bernardo D, Motoi H, Sonoda M, Kuroda N, et al. Scalp EEG interictal high frequency oscillations as an objective biomarker of infantile spasms. Clin Neurophysiol [Internet]. 2020;131(11):2527–36. Available from: https://doi.org/10.1016/j.clinph.2020.08.013

Nehlig A, Coppola A, Moshe S. Epileptic Syndromes in Infancy, Childhood, and Adolescence. 5th ed. Bureau M, Genton P, Dravet C, Delgado-Escueta AV, Tassinari CA, Thomas P, et al., editors. Montrouge, France: John Libbey Eurotext; 2012.

O’Callaghan FJK, Lux AL, Darke K, Edwards SW, Hancock E, Johnson AL, et al. The effect of lead time to treatment and of age of onset on developmental outcome at 4 years in infantile spasms: Evidence from the United Kingdom Infantile Spasms Study. Epilepsia. 2011;52(7):1359–64.

Osborne JP, Lux AL, Edwards SW, Hancock E, Johnson AL, Kennedy CR, et al. The underlying etiology of infantile spasms (West syndrome): Information from the United Kingdom Infantile Spasms Study (UKISS) on contemporary causes and their classification. Epilepsia. 2010;51(10):2168–74.

Pavone P, Striano P, Falsaperla R, Pavone L, Ruggieri M. Infantile spasms syndrome, West syndrome and related phenotypes: What we know in 2013. Brain Dev. 2013;36(9):739–51.

Peng CK, Buldyrev S V., Havlin S, Simons M, Stanley HE, Goldberger AL. Mosaic organization of DNA nucleotides. Phys Rev E. 1994;49(2):1685–9.

Peng CK, Buldyrev SV, Goldberger AL, Havlin S, Sciortine F, Simons M, et al. Long-range correlations in nucleotide sequences. Nature. 1992;356(6365):168–70.

Primec ZR, Stare J, Neubauer D. The risk of lower mental outcome in infantile spasms increases after three weeks of hypsarrhythmia duration. Epilepsia. 2006;47(12):2202–5.

Van Putten MJAM, Stam CJ. Is the EEG really “chaotic” in hypsarrhythmia? IEEE Eng Med Biol Mag. 2001;20(5):72–9.

Riedl M, Muller A, Wessel N. Practical considerations of permutation entropy: A tutorial review. Eur Phys J Spec Top. 2013;222(2):249–62.

Riikonen RS. Favourable prognostic factors with infantile spasms. Eur J Paediatr Neurol [Internet]. 2010;14(1):13–8. Available from: http://dx.doi.org/10.1016/j.ejpn.2009.03.004

Rosso OA, Hyslop W, Gerlach R, Smith RLL, Rostas JAP, Hunter M. Quantitative EEG analysis of the maturational changes associated with childhood absence epilepsy. Phys A Stat Mech its Appl. 2005;356(1):184–9.

Schwender D, Daunderer M, Mulzer S, Klasing S, Finsterer U, Peter K. Spectral edge frequency of the electroencephalogram to monitor “depth” of anaesthesia with isoflurane or propofol. Br J Anaesth. 1996;77(2):179–84.

Shannon CE. A Mathematical Theory of Communication. Bell Syst Tech J [Internet]. 1948;5(3):3. Available from: http://portal.acm.org/citation.cfm?doid=584091.584093%5Cnhttp://ieeexplore.ieee.org/lpdocs/epic03/wrapper.htm?arnumber=6773024

Shields WD. Diagnosis of infantile spasms, Lennox-Gastaut syndrome, and progressive myoclonic epilepsy. Epilepsia. 2004;45 Suppl 5:2–4.

Shrey DW, Kim McManus O, Rajaraman R, Ombao H, Hussain SA, Lopour BA. Strength and stability of EEG functional connectivity predict treatment response in infants with epileptic spasms. Clin Neurophysiol [Internet]. 2018;129(10):2137–48. Available from: https://doi.org/10.1016/j.clinph.2018.07.017

Siniatchkin M, Van Baalen A, Jacobs J, Moeller F, Moehring J, Boor R, et al. Different neuronal networks are associated with spikes and slow activity in hypsarrhythmia. Epilepsia. 2007;48(12):2312–21.

Smith RJ, Shrey DW, Hussain SA, Lopour B. Quantitative Characteristics of Hypsarrhythmia in Infantile Spasms. Conf Proc IEEE Eng Med Biol Soc. 2018;538–41.

Smith RJ, Sugijoto A, Rismanchi N, Hussain SA, Shrey DW, Lopour BA. Long-Range Temporal Correlations Reflect Treatment Response in the Electroencephalogram of Patients with Infantile Spasms. Brain Topogr. 2017;30(6).

Stam CJ, Nolte G, Daffertshofer A. Phase Lag Index: Assessment of Functional Connectivity From Multi Channel EEG and MEG with Diminished Bias from Common Sources. Hum Brain Mapp. 2007;28(11):1178–93.

Stamps F, Gibbs E, Rosenthal I, Gibbs F. Treatment of Hypsarrhythmia with ACTH. JAMA. 1959;171(4):116–9.

Sue WC, Mikati MA, Kramer U. Hypsarrhythmia: Frequency and variant patterns and correlation with etiology and outcome. Neurology. 1997;48(1):197–203.

Widjaja E, Go C, McCoy B, Snead OC. Neurodevelopmental outcome of infantile spasms: A systematic review and meta-analysis. Epilepsy Res [Internet]. 2015;109:155–62. Available from: http://dx.doi.org/10.1016/j.eplepsyres.2014.11.012

Wu JY, Koh S, Sankar R, Mathern GW. Paroxysmal Fast Activity: An Interictal Scalp EEG Marker of Epileptogenesis in Children. Epilepsy Res. 2008;82(1):99–106.

